# High Temporal Resolution Measurements of Movement Reveal Novel Early-Life Physiological Decline in *C. elegans*

**DOI:** 10.1101/2021.09.07.459324

**Authors:** Drew B. Sinha, Zachary S. Pincus

## Abstract

Age-related physiological changes are most notable and best-studied late in life, while the nature of aging in early- or middle-aged individuals has not been explored as thoroughly. In *C. elegans*, studies of movement *vs.* age generally delineate three distinct phases: sustained, youthful movement; a discrete onset of rapidly progressing impairment; and gross immobility. We investigated whether this first period of early-life adult movement is simply a sustained “healthy” level of high function followed by a discrete “movement catastrophe” — or whether there are early-life changes in movement that precede future physiological declines. To determine how movement varies during early adult life, we followed isolated individuals throughout life with a previously unachieved combination of duration and temporal resolution. By tracking individuals across the first six days of adulthood, we observed declines in movement starting as early as the first two days of adult life, as well as high interindividual variability in total daily movement. These findings suggest that movement is a highly dynamic behavior early in life, and that factors driving movement decline may begin acting as early as the first day of adulthood. Using simulation studies based on acquired data, we suggest that too infrequent sampling in common movement assays limits observation of early-adult changes in motility, and we propose feasible alternate strategies and a framework for designing assays with increased sensitivity for early movement declines.

## Introduction

Of many physiological changes that occurs during aging, loss of mobility is a critical factor in loss of health and well-being. Changes in mobility in humans are associated with changes in other functional domains (e.g. cognition[1,2]), and decreased mobility predicts future mortality in both humans[3,4] and model organisms[5,6]. Given the importance of mobility and the ease of measuring mobility during aging in humans and model organisms, considerable effort has gone into the study of mobility throughout life.

Similar to humans, the model organism *C. elegans* shows stereotypic age-related changes in movement and neuromuscular function. This makes it a very tractable model with which to study the timecourse of age-related changes in mobility. Physiological processes seen in humans have also been implicated in movement decline of *C. elegans*, such as sarcopenia[5,7] and age-related deterioration of neurons[8–10].

Previous work suggests that age-related changes in *C. elegans* movement follow a stereotyped pattern: a period of consistently high, vigorous activity in early life (approximately days 1-6 of adulthood at 20°C); followed by rapid decline and loss of spontaneous movement; and eventually gross immobility even to external stimulus[11–15]. However, these observations of a relatively uneventful, unchanging early adulthood stand in contrast to what is known of other early adult *C. elegans* physiology. Indeed, early adulthood is a time of otherwise great change, including: continued changes in size; transition into and out of reproductive maturity[16]; changes in food consumption[17] and defecation[18]; and increased propensity for extrinsic damage from environmental factors such as colonization by bacterial food[19]. This discrepancy raises a straightforward question: does movement truly remain at a constant level of “healthy” function throughout early adulthood? Or is movement in these young animals subject to “aging” as well?

One plausible reason why early-life movement in these animals appears unchanging is that movement assays are not optimized to discern between different levels of vigorous activity. On standard vermiculture plates, animals can travel at high speeds (up to ~500 μm/s[20]), making observation of movement challenging. A number of strategies/technologies have emerged to track animals, ranging from simple video imaging or custom tools[21,22] for tracking animals on plates to novel culture systems with specialized imaging systems[12,23]. In many aging studies, especially longitudinal studies using custom systems/software, investigators typically pursue one of two strategies: obtain either frequent, low-quality measurements of movement (a handful of images obtained every few hours)[11]; or infrequent, high-quality movement measurements (e.g. once-daily videos of several minutes)[12–14,24]. The latter strategy has historically been generally more common. It is not clear, however, the degree to which movement measures from either of these two strategies correlate with “perfect” (if unobtainable) observations of movement: continuous, lifelong video-recording of every individual. As such, it is also not clear whether the failure to observe changes in early-life movement in *C. elegans* reflects aging biology or technical limitations.

We thus investigated whether young-adult animals experience age-related declines in movement by collecting movement data at previously unattained spatial and temporal resolution. To measure longitudinal changes in animal movement at high temporal resolution, we employed the *C. elegans* Worm Corral longitudinal culture system[11,25]. This system has the salient limitation that each individual is confined to a relatively small area (a circular pad of bacterial food ~1.5-2 mm in diameter). However, this allows us to efficiently make repeated measurement of changes in behavior for multiple, isolated individuals across multiple days. Using this system, we demonstrate that second-scale animal movement is surprisingly well-represented by measurements of animal movement at considerably lower temporal resolution (e.g. 1 frame per minute; fpm). That established, we tracked movement of several individuals at high fidelity with minute-to-minute measurements over the course of several days of early adult life. From these data, we found substantial decline and interindividual variability occurring during this period of life – which is usually thought of as uniform, vigorous movement. Finally, using simulations based on the minute-to-minute data acquired in this study, we suggest that typical protocols used to assess animal movement in standard vermiculture may be insensitive to early-life physiological changes. To address this concern, we propose several simple improvements to increase sensitivity to age-related movement changes. Overall, this study not only suggests the presence of important age-related physiological changes in mobility in early adulthood, but provides a schema by which investigators can verify the fidelity of data-acquisition strategies and their sensitivity to early-life mobility changes.

## Methods

### Animal Husbandry

For reasons unrelated to this analysis, all experiments in this study use the *C. elegans* strain BA671 [*spe-9(hc88)*]. This strain has a temperature-sensitive fertility defect above 25°C, but is otherwise phenotypically wild type at 25°C and completely phenotypically wild-type at 20°C[26]. Strains were maintained at 20°C on standard NGM plates seeded with *E. coli* OP50[27].

### Worm Corral Bacterial Food & Animal Preparation

We maintained our lifelong culture system at 20°C for these experiments to ensure the movement results would be as comparable as possible to standard assays which are also typically performed at 20°C. As *spe-9(hc88)* individuals are fertile at this temperature, we prevented progeny accumulation by instead employing feeding RNAi using the RNAi-compatible OP50-derivative strain OP50(*xu363*)[28] transformed to produce dsRNA against the embryonic transcription factor *pos-1*. Animals fed *xu363/pos-*1 during late development and early adult life produce fertilized, but developmentally arrested eggs.

Concentrated *xu363/pos-1* bacterial food for Worm Corrals was prepared as follows. First, a culture was prepared by inoculating LB with a single colony of *xu363/pos-1* and incubated at 37°C for 17 hours (“overnight”). This overnight culture was diluted down to an optical density (OD) of 0.25, grown up to an OD of 0.5, and induced with addition of isopropyl beta-d-1-thiogalactopyranoside (IPTG) at 1 mM final concentration followed by further incubation for 3 hours. The bacterial culture was resuspended in M9 salts solution at ~75% wt/vol and stored at 4°C for no more than 3 days.

Embryos were prepared for use in Worm Corrals by setting up P0 animals for a synchronized egg-lay. P0s were synchronized at the L4 stage and, the following day, were transferred to a new plate to lay eggs. To prevent cross-contamination between *xu363/pos-1* and OP50 (which doubles faster and would thus otherwise outgrow the *xu363/pos-1*), the egg-lay plate was seeded with a control *xu363* strain transformed to produce dsRNA against GFP, and transferred animals were deposited off the *xu363/GFP* lawn. Embryos were picked from the egg-lay plate 17-18 hours after transfer of P0s. For each experiment, embryos were picked from a single plate to the worm corral device at room temperature, and were out no more than 1-1.5 hours prior to initiation of experiment.

### Construction of Longitudinal Culturing Devices

For longitudinal observation experiments, custom solid media longitudinal culturing devices ( “Worm Corrals”) were used[11,25]. Briefly, the solid media in these devices consists of a PEG-based hydrogel containing a modified formulation of NGM (pH 6.3 with 20 μg/mL cholesterol in solution and lacking calcium chloride). The mixed solution was pipetted in a liquid state into a reservoir formed from a 2.5 mm thick aluminum frame affixed to a standard microscope glass slide, and left to cross-link for 1.5 hrs at room temperature. After crosslinking, 0.5 μl droplets of concentrated *xu363/pos-1* bacteria were pipetted onto the hydrogel surface, and a single embryo was transferred from an egg-lay plate into the drop manually with an eyelash to minimize disturbance of the food pad. After loading the hydrogel with bacteria & embryos, the device was sealed by adding a thin layer of polydimethylsiloxane (10:1 base:cure ratio) onto the hydrogel surface.

Devices were placed on the stage of an automated compound microscope during subsequent culturing and observation. The stage was housed in a sealed enclosure that was kept at constant temperature (20°C) and humidity (> 90% relative humidity) throughout the duration of the experiment.

### Data Acquisition

During all experiments, standard brightfield imaging was performed with a compound microscope (5× objective) fitted with a Zyla 5.5 sCMOS camera (sensor size 16.6 x 14 mm; pixel size 6.5 μm; Andor Instruments, Belfast, UK). Prior to data collection, animals were allowed to progress through larval development for 48 hours, such that data acquisition began just prior to the beginning of adulthood for all animals. During this initial period, individuals were followed by taking a single brightfield image every three hours to assess for proper development.

For experiments in which data was collected at 1 frame per second (fps), each individual was continuously imaged in standard brightfield conditions for a total of 20 minutes (allowing for a theoretical maximum of 72 animals to be serially imaged during a 24 hour period). For experiments in which data was collected at 1 frame per minute (fpm), all animals were imaged at each timepoint. In these latter experiments, movement was also stimulated every 3 hours by exposure to a blue LED lamp (Spectra light engine; Lumencor Beaverton, OR; 470 nm center frequency) for 500 ms. However, because this study is restricted to unstimulated movement, data from the 30 minutes following each stimulation were excluded from further analysis.

### Post-Experimental Image Processing, Data Analysis, and Data Simulation

The location of each animal in every image was identified using a convolutional neural network, as in previous work[29] (Fig 1A). To assess consistency of bacterial food pad sizes, the bacterial lawn was identified by fitting a two-mode Gaussian mixture model to the image intensity histogram, and taking all pixels corresponding to the lower-intensity mode as associated with the (visually darker) lawn. This procedure was performed for the first five images of that image series, and these binary image masks were combined with a bitwise OR operation to obtain a robust estimate of the entire lawn region (Fig 1A-B). These masks were verified by manual inspection. Onset of adulthood for each animal was identified manually by an experimenter and all reported times are specified with respect to the first egg laid, 3-6 hours after the exit from the L4 molt[11]. In this study, movement was measured as the change in position of the centroid of the animal across two or more images.

**Fig 1:**
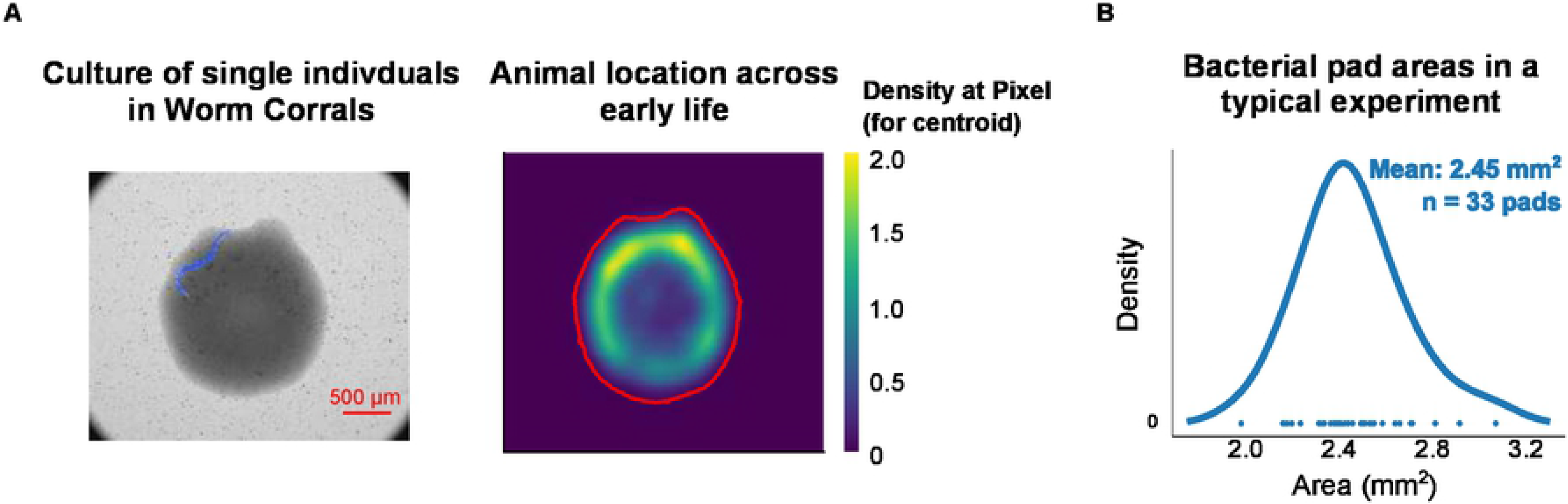
Experimental setup and correlation between second- and minute-scale movement. (A) *Left panel*, Imaging of single individuals in Worm Corrals were performed at regular intervals using standard brightfield imaging on a compound microscope. In each image, the region corresponding to a worm (blue region) and food pad were identified. *Right panel*, the animal’s location over the first six days of adulthood represented using a 2-dimensional kernel density plot of the centroid of the worm region. Most animals tended to spend most timepoints near the edge of the bacterial food pad where, due to the “coffee ring effect”[31], bacterial density is greatest. (B) Distribution of lawn area measurements from segmentation. The mean lawn area was ~2.5 mm^2^ (equivalent to a circle with a radius of 0.88 mm/diameter of 1.76 mm).

*P*-values for Pearson linear regression/correlation coefficients were calculated using the standard F-test. We defined a correlation as of biological interest if it reached statistical significance at *p* < 0.05 (corrected for multiple comparisons by Bonferroni correction as appropriate) and yielded a coefficient of determination (R^2^) > 0.10. Correlations with lower coefficient of determinations, regardless of *p*-value, do not capture a great deal of the relevant biological effect and empirically have proven difficult to reproduce.

Simulated timecourses of movement were generated by subsampling data acquired at 1 fpm. To sample the “Single Snapshot” and the “Hybrid Movement” protocols, individual images/worm positions were taken at the prescribed sampling interval from the data. To simulate the “Daily Video” protocol (15 minutes of images obtained at 1 fps), we simply used 15 minutes of images obtained at 1 fpm, from the beginning of each 24-hour period. Though these two recording regimes seem very dissimilar, we found from our small-scale 1 fps pilot dataset (Fig 2) that there is a strong correlation between total displacement over 15 minutes at 1 fps and net displacement at (subsampled) 1 fpm imaging (R^2^ = 0.81; S1 Fig).

**Fig 2:**
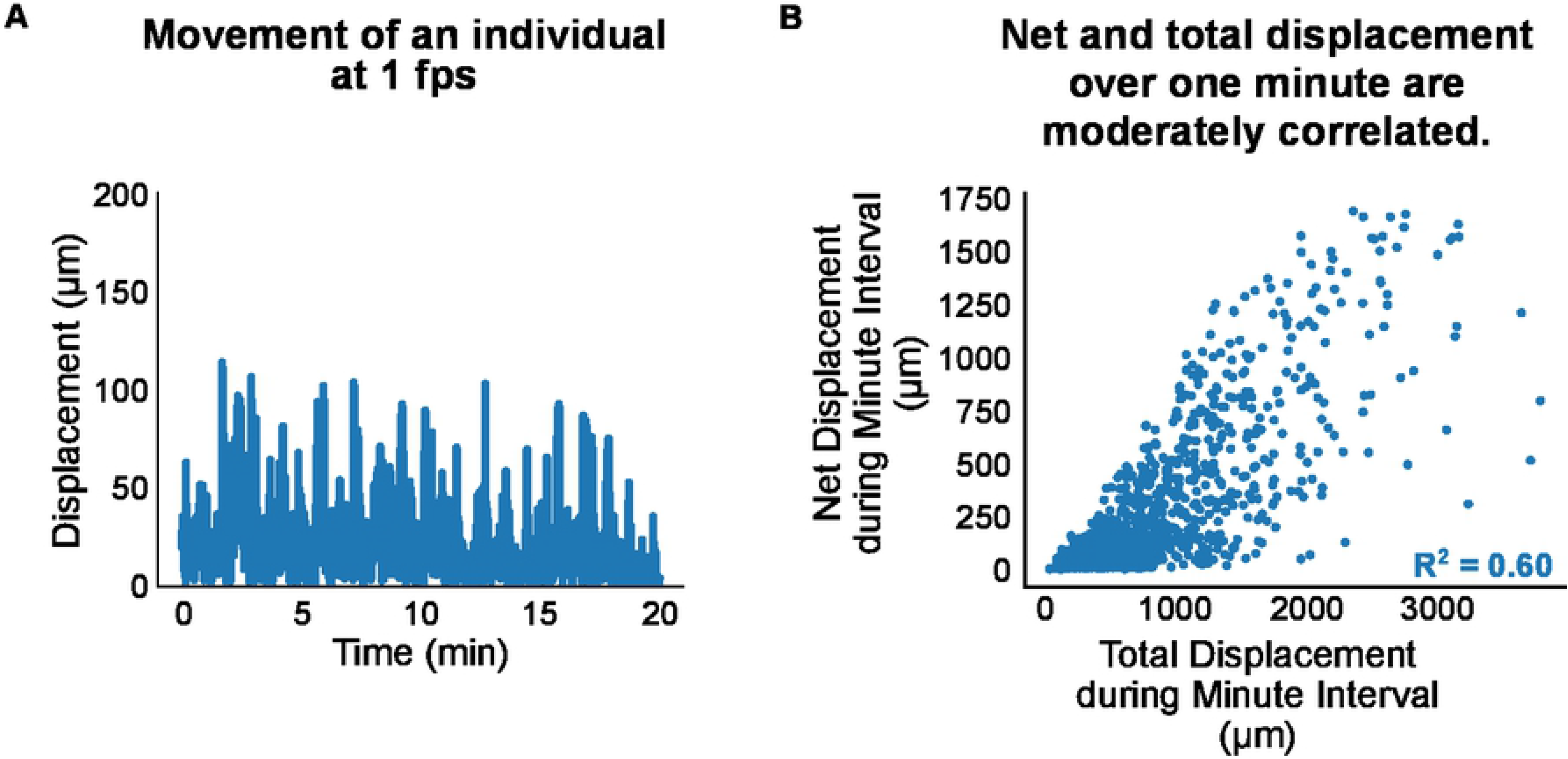
Experimental setup and correlation between second- and minute-scale movement. (A) Representative movement activity (inter-timepoint centroid displacement) of a day 1 adult animal assessed at 1 fps for 20 minutes. (B) Comparison of total displacement across a one minute interval at 1 fps vs net displacement over the same interval at 1 fpm over the same interval (n = 48 animals * 20 minute intervals = 9600 data points). The overall correlation between net and total displacement is strong with an R^2^ = 0.60 (95% CI 0.50-0.68 generated by bootstrap resampling measurements of random subsets of 43 animals/90% of animals).

## Results

### Minute-scale measurements of movement correlate with finer animal movement

To investigate the movement behavior of early-adult animals, we first set out to determine minimal conditions for reliably measuring movement in young animals over multiple days. Ideally, each animal under study would be continuously imaged throughout the experiment at a frame rate and magnification sufficiently high to capture all motion. This is of course generally impractical, as magnification, frame rate, number of animals studied, and frequency of imaging each individual all generally trade off against one another. These trade-offs led us to consider the degree to which an animal’s second-scale movement is well-represented by measurements taken at longer intervals.

We assessed whether minute-to-minute measurements were sufficient to represent finer movement activity. We serially acquired 20-minute video recordings (at 1 fps) of day 1 adult individuals, isolated from one another via the Worm Corral system (Fig 1 & Fig 2A). Based on previous studies, video imaging at 1 fps is generally sufficient for measuring forward movement, though likely captures only part of other major movement behaviors (particularly intermittent reverse movements – so called “reversals” – whose duration is on the order of 1-2 seconds or less)[20,30]. For a given minute during the recording session, we compared the total distance traveled by an animal (at 1fps) to its net displacement across the first and last images taken over that minute (i.e. at an effective 1 fpm; Fig 2A). We observed a strong correlation between second- and minute-scale movement for any given minute interval of the session (mean R^2^ = 0.60, bootstrap 95% CI 0.50-0.68; Fig 2B). These findings suggest that, on average, 1 fpm recordings of displacement capture over half of the inter-individual variance present in second-scale movement recordings.

### Minute-scale recordings of early-adult movement reveal early-life movement decline

Because minute-scale recordings capture an appreciable fraction of total movement, we used these measures to determine how movement varies during the first days of early adulthood in *C. elegans*. The first 4-6 days of adulthood are typically known as a period of vigorous activity prior to the beginning of an overt, qualitative decline in movement rates[7,11–13,32]. Consequently, we chose to focus on this period when assessing patterns of early-adult decline.

We collected movement data from 33 individuals at 1 frame per minute (fpm) throughout the first 6 days of adulthood. Consistent with prior reports, there were substantial within-individual variability in minute-to-minute; additionally, we observed substantial interindividual variability in minute-to-minute animal movement. Some individuals showed relatively high and/or sustained periods of maximal movement (Fig 3A, *left panel*) while others exhibited a relatively lower baseline level of activity (Fig 3A, *right panel*).

**Fig 3:**
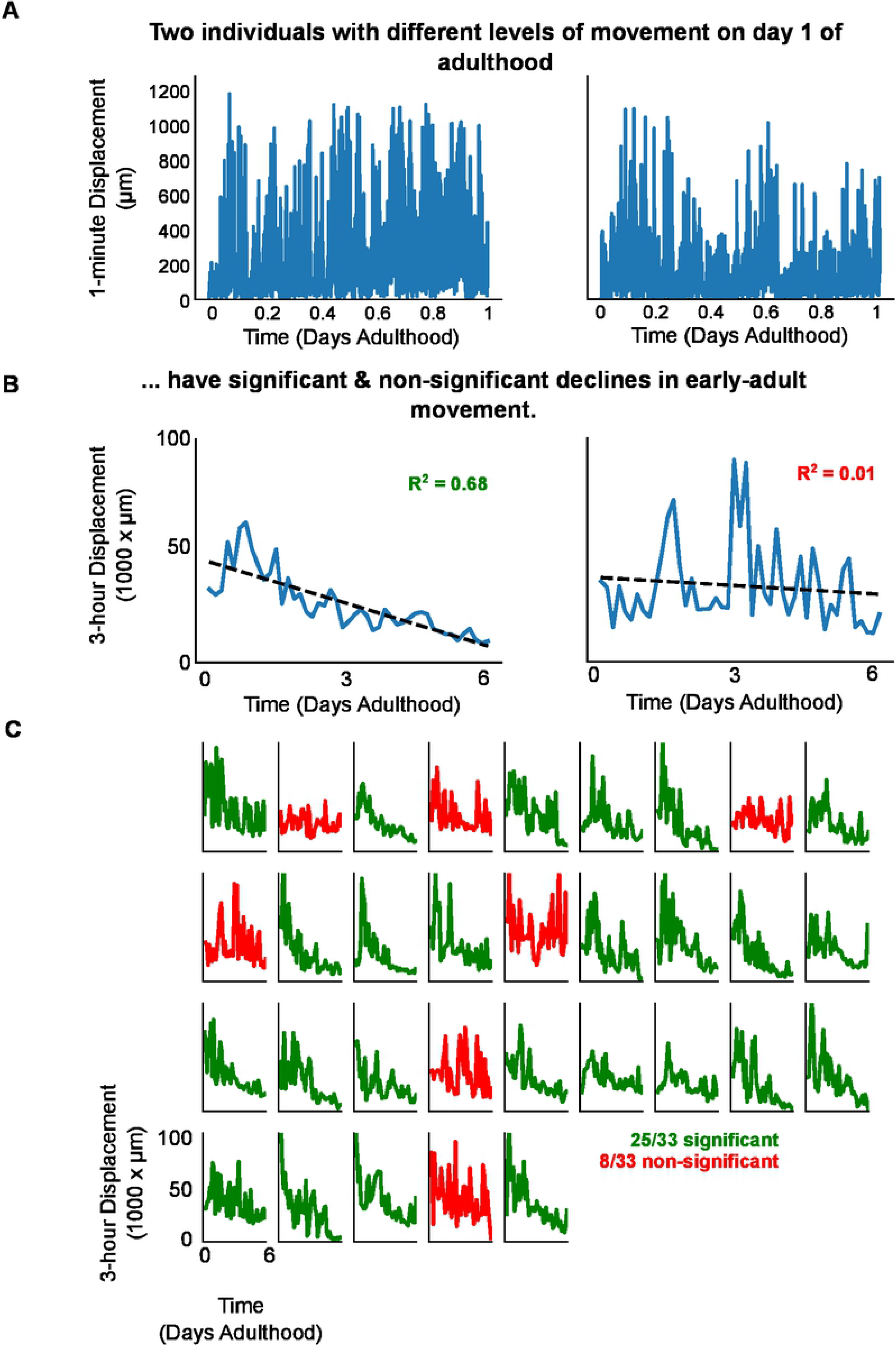
Animal movement shows substantial decline during early-adult life. (A) Examples of movement taken at 1 fpm during the first day of life. Shown are data from two individuals with relatively high (*left panel*) and low (*right panel*) movement. (B) Corresponding movement declines for the respective individuals in (A), integrated over 3-hour intervals to yield 8 timepoints a day. The left-panel individual’s activity has a significant decline whereas the right-panel individual’s does not. (C) Visual depiction of all 33 individuals’ activity during the first six days of life, colored by significant (green) vs. non-significant (red) declines over time.

We next examined how each individual’s level of activity changes during early adulthood. We quantified total movement at 3 hour intervals by summing the total minute-to-minute displacement over each 3 hour interval (Fig 3B)[11,33]. On this time scale, we observed substantial decline in movement for most animals: 26/33 animals (~79%) showed a statistically significant decline in movement across the first six days of adulthood (Fig 3C). Overall, these findings suggest that at least some animals do experience substantial declines in movement during the first days of adult life.

### Early adult movement is variable between individuals and is not a persistent trait

The observation that movement behavior can decline during early adulthood raises a related question – do all individuals experience a similar pattern of decline? More specifically, how different are individuals’ movement behavior across the first few days of adulthood? On cursory inspection, animals show substantial differences in daily activity even on the first day of adulthood (Figs 3A & B). When animal movement was quantified by total minute-to-minute distance traveled during day 1 of adulthood, total movement spanned a 4-fold range across all animals (Fig 4A). Expressed as a coefficient of variation (a normalized ratio of the standard deviation to mean of total day movement; CV), this level of variability amounts to ~1/4 (27%) of the average distance that an animal traveled that day. Interestingly, this variability was evident across each day and increased during days 5 & 6 as animal movement continued to decline (Fig 4A).

**Fig 4:**
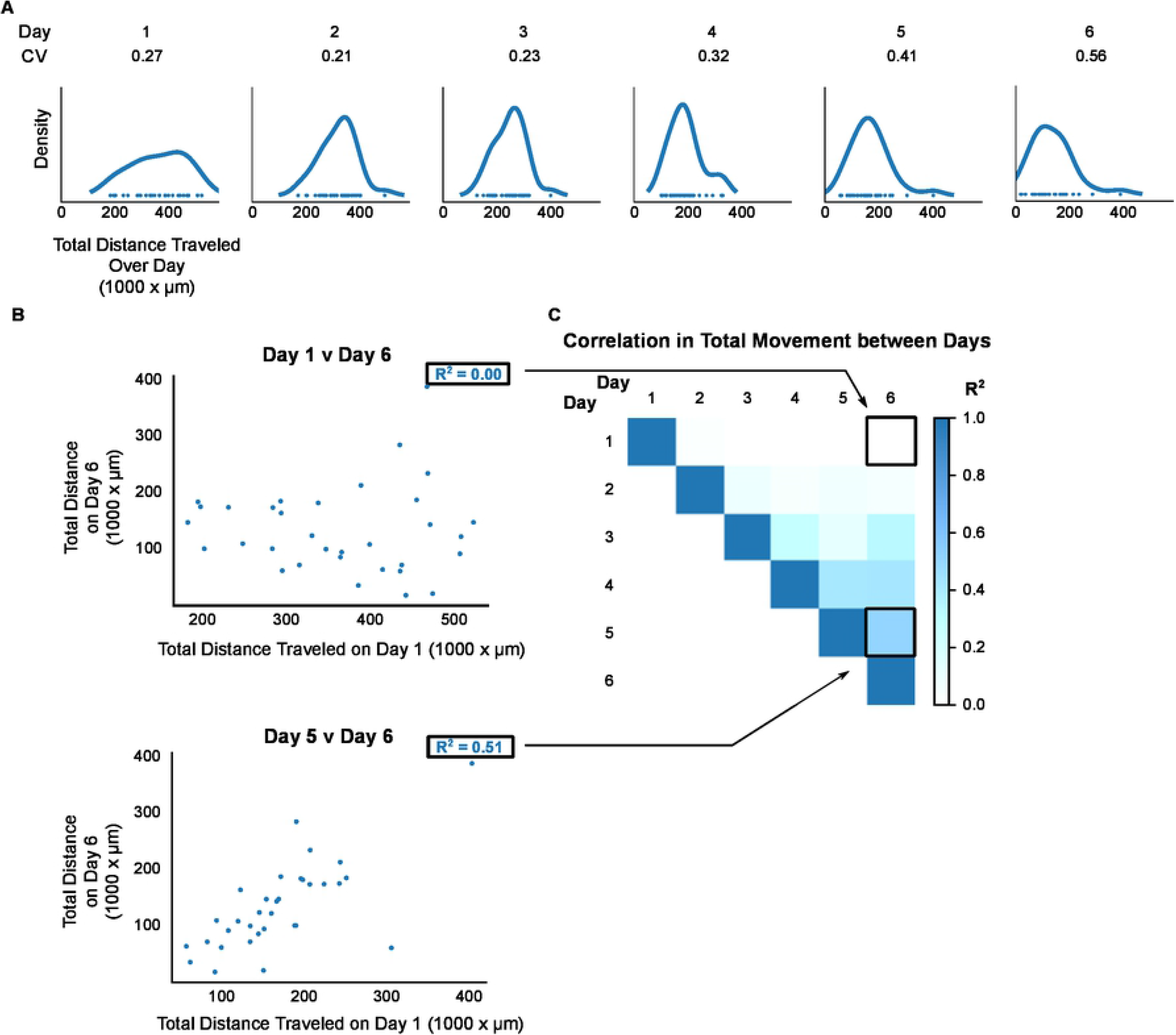
Interindividual variability and persistence of movement activity throughout early-adult life. (A) Distribution of total movement on each day of the first 6 days of adulthood across all observed individuals. The population is depicted as a kernel density histogram with each dot at the bottom of the histogram representing a single individual’s movement for the day. Variability in movement on a given day is quantified as coefficient of variation (CV) and shown above each histogram. (B) Scatter plot demonstrating degree of correlation between an individual’s first- (*top panel*) or fifth-day (*bottom panel*) movement and its movement on the sixth/last day of observation. Correlation noted as Pearson-product linear R^2^. (C) Heatmap of pair-wise correlations in movement between all days of observation. Entries corresponding to each panel in (B) depicted by arrows.

Having observed such variability across animals in the setting of significant age-related declines, we questioned whether an animal’s level of movement reflected a persistent trait; that is, do some animals always move more than others, even in the midst of an overall decline in movement with age? We thus asked if movement on any given day was predictive of movement on a future day. Surprisingly, we found that movement among the first 2-3 days of adulthood was not predictive of movement on any other day out to the end of early-adulthood on day 6 (Fig 4B, *top panel*; Fig 4C). Greater correlations among daily movement scores began to emerge at days 4-5 of adulthood (Fig 4B, *bottom panel*; Fig 4C). Taken together, these findings suggest that youthful movement is highly variable both between and within individual animals, despite being overlaid on an overall downward trend through early adulthood.

### Standard movement assays are underpowered to resolve early adult movement decline

It is curious that, despite extensive study of *C. elegans* movement through aging, early adulthood movement declines have not previously been noted. One reason for this is likely the high degree of between- and within-individual variability in movement noted above. However, we also wondered whether the design of specific data-acquisition protocols could exacerbate this issue. To answer this question, we performed simulations to study the effects of different measurement protocols on the types of conclusions that can subsequently be drawn.

In these simulation studies, we resampled image data from the collected “gold-standard” data collected at 1 fpm and to simulate the movement scores that would be generated for each individual under different data-collection protocols. For consistency with the analyses of minute-to-minute movement in previous sections, we aggregate movement into 3-hour timepoints when comparing protocols. We then compared the resulting series of reconstructed movement measurements to those calculated from the original data.

We examined two different measurement protocols, both common in the *C. elegans* literature (Fig 5A). First, we defined a “Single Snapshot” protocol that measures movement as the displacement of an animal between timepoints at a prescribed interval (i.e. displacement in animal position between two images taken 2 minutes to almost 3 hours apart)[11]. Second, we defined a “Daily Video” protocol that measures 15 minutes of movement at 1 fps, once a day per individual. This is analogous to standard plate assays in which animals are recorded at high frame rate for a fixed amount of time each day[14,23,32,34]. Unlike the “Single Snapshot” protocol which we can readily examine by subsampling our continuous 1 fpm imaging data, we cannot directly produce a dataset equivalent to the “Daily Video” protocol from the images obtained. However, we noted from down-sampling our 1 fps pilot data to 1 fpm that summed minute-to-minute movement across a 15-minute interval is very strongly correlated with summed second-to-second movement over that same interval (R^2^ = 0.81; see S1 Fig). Thus, we can effectively simulate “Daily Video” measurements for a given day by summing minute-to-minute displacement over 15 minutes at the start of that 24 hour period.

**Fig 5:**
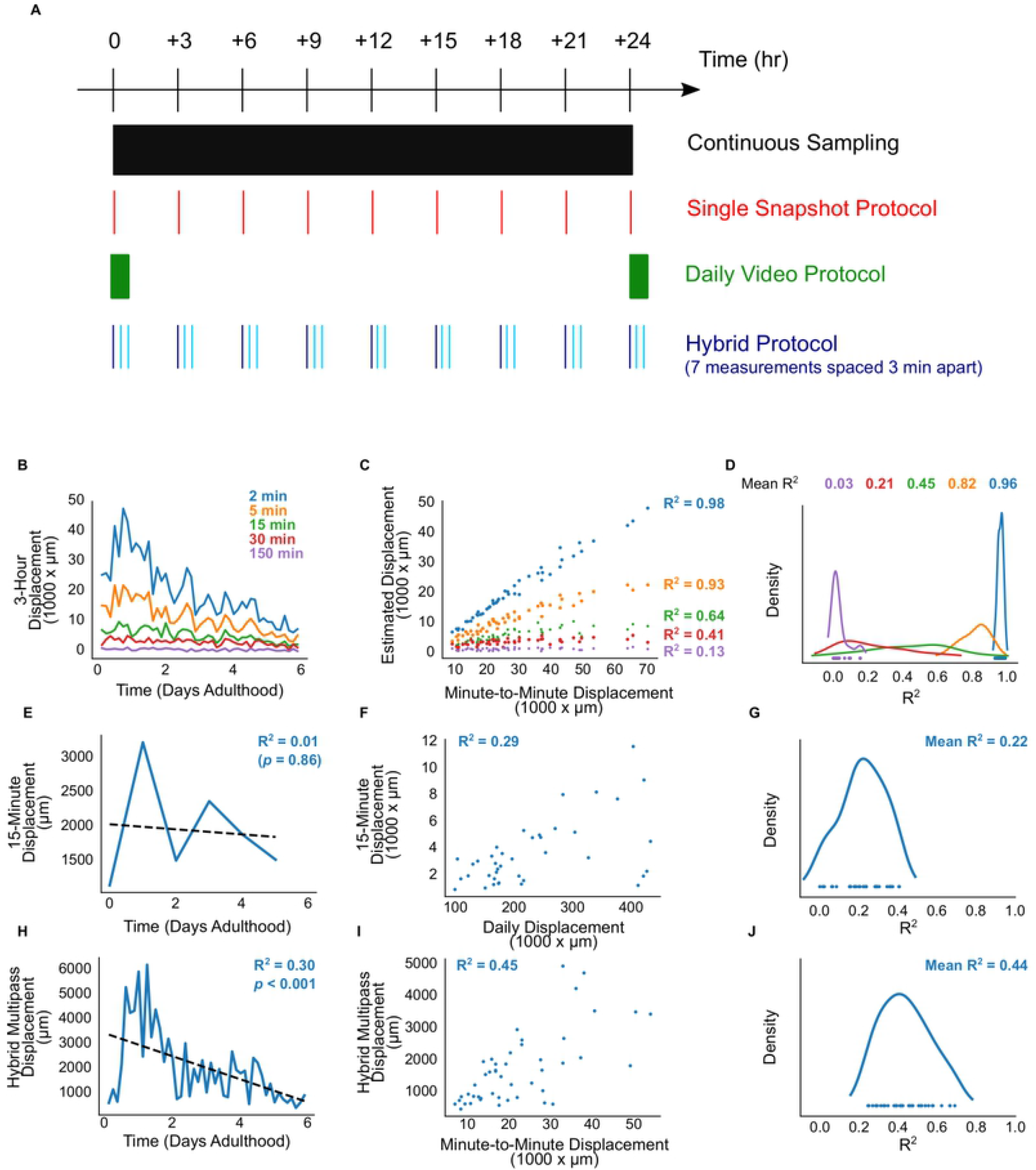
Comparison of validity of three movement protocol schemes. (A) Graphical cartoon of the various protocols of interest. The “Continuous Sampling” condition is the baseline-quality data collected in this study. In the “Single Snapshot” protocol, one image per timepoint is taken, and the movement score for that timepoint is the displacement of the individual’s centroid from the previous timepoint. The “Daily Video” protocol consists of 15 images obtained per timepoint at 1 fpm; the movement score for that timepoint is the summed total displacement across all 15 images acquired in that day’s timepoint. Finally, in the “Hybrid Multipass” protocol, several images per timepoint are taken with a wider temporal spacing (allowing image acquisition for several individuals to be interleaved) and the total displacement across those images is used as the movement score for that timepoint. Here, we acquire 7 images each spaced 3 minutes apart (for a total interval of 18 minutes). (B-D) Illustration of data generated by the Single Snapshot protocol at various inter-timepoint intervals. (B) One individual’s activity across the first 6 days of adulthood. (C) For that individual, the correlation between the total displacement during those intervals (measured from gold-standard 1 fpm data) and net displacement using the specified inter-timepoint interval. (D) Kernel density plot showing the distribution of correlations between 1fps and Single Snapshot obtained for all individuals at the various inter-timepoint intervals. (E-G) Illustration of the data generated by the Daily Video protocol at various inter-timepoint intervals. (E) Activity from the same individual in panel B, across the first 6 days of adulthood. (F) The correlation between the total displacement during the sampling period (from 1 fpm data) and that day’s movement score from the Daily Video data for this individual. (G) Kernel density plot showing the distribution of correlations between gold-standard and Daily Video data, across all individuals. (H-J) Illustration of data generated using the Hybrid Multipass protocol. (H) Activity from the same individual in panel B, across the first 6 days of adulthood. (I) The correlation between the total displacment during (with 1 fps data) and the Hybrid Multipass movement score for this individual. (J) Kernel density plot showing the distribution of correlations between gold-standard and Hybrid Multipass data, across all individuals.

We found that the Single Snapshot protocol was unable to recapitulate decline in animals seen with 1 fpm imaging. With increasing duration between subsequent images, an animal’s displacement between images more poorly resembles total 1 fpm movement over time (Fig 5B). Across all timepoints during early life for a given animal, Single Snapshot movement correlates increasingly worse with overall movement as the inter-image interval is increased (Fig 5C). Across all animals, this correlation falls to a moderate correlation at 15-minute inter-image intervals (mean R^2^ = 0.44), and virtually no correlation at 150 minute intervals (mean R^2^ = 0.03; Fig 5D). At this last inter-image interval, the Single Snapshot protocol does not identify age-related movement decline in any individual (Fig 6). Taken together, these findings suggest that inter-image intervals of 2-5 minutes are most representative of an animal’s overall movement across the lifespan.

**Fig 6:**
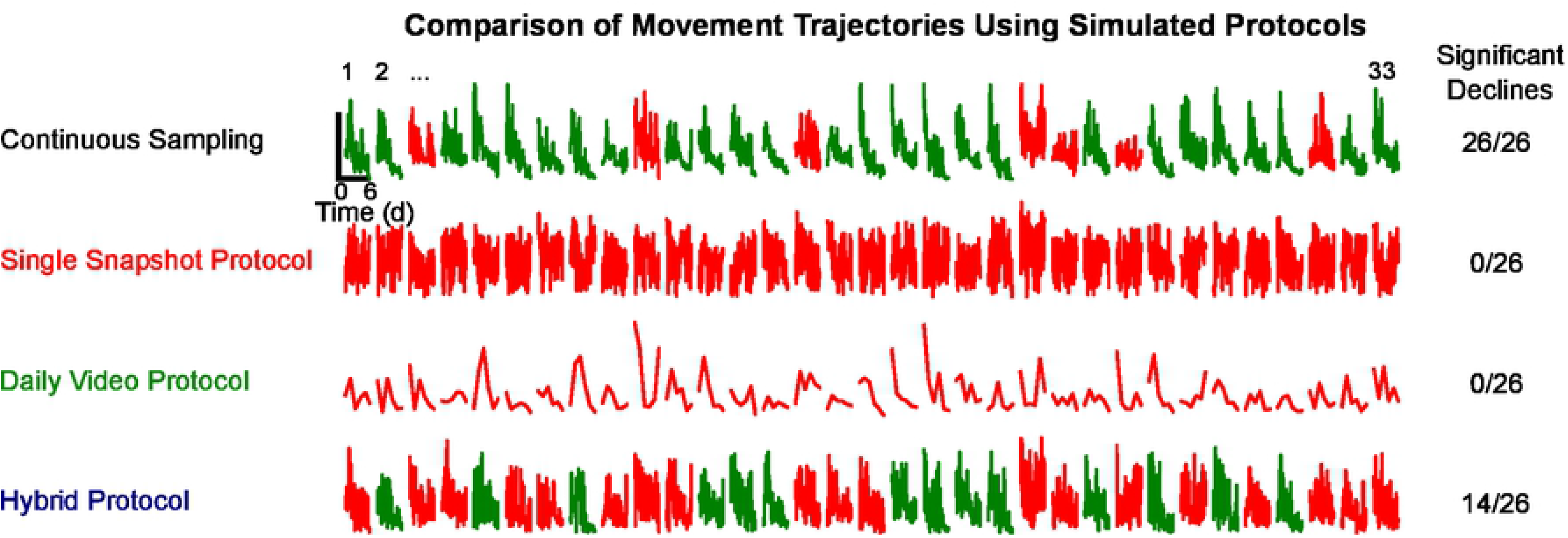
Visual comparison of measurable decline in activity across different image acquisition protocols. Activity traces are labeled for all 33 animals and colored red vs. green to denote non-significant vs. significant decline, respectively (as in Fig 3C).

The Daily Video protocol had similar difficulty in recapitulating overall movement across this period. Qualitatively, an animal’s total summed movement across a whole day is unlikely to be well-approximated by a single short burst of observations (Fig 5E), unless movement across all fifteen-minute intervals throughout the day are very highly correlated. Reports of periodic/diurnal patterns in *C. elegans* movement behavior[35–37] make this *a priori* unlikely. And indeed, movement during the specified 15-minute interval correlates at best modestly with each day’s total movement (mean R^2^ across animals = 0.22; Fig 5F-G). Moreover, when used to assess early life movement decline, the Daily Video protocol fails to identify age-related movement decline in any individual (Fig 6). Of note, assessing movement via 15-minute samples twice per day (as done in some studies) yielded a improved correlation between 15-minute movement and (half-day) movement activity (mean R^2^ across animals = 0.46), but did not affect these conclusions (S2 Fig).

### Optimization of measurement protocols produces a simplified assay with improved sensitivity for early-adult movement decline

These results suggest that maximizing the quantity of measurements (number of timepoints) or quality of measurements (data acquisition per timepoint) individually fails to reproduce trends in early-life movement seen with continuous (1 fpm) image acquisition. This caused us to wonder whether a protocol with measurements of intermediate frequency and quality might fare better.

We thus evaluated a “Hybrid Measurement” protocol that combines sparser imaging over a relatively long duration of acquisition. This protocol was motivated by the observation that movement on the order of a few (1-6 minutes) minutes is well-correlated to second-scale movement (e.g. Fig 5B). As a proof of principle, we used this observation to maximize the quality of measurements for our group’s Worm Corral system. Specifically in our workflow, separately housed individuals on up to 6-7 devices are serially imaged on an upright compound microscope at high-magnification with a typical throughput of ~600-700 animals with a ~3 hour inter-timepoint interval. On average, moving to a new individual on a device, automatically focusing, and acquiring an image takes ~1.4 sec. Under these constraints, we devised a protocol in which movement is measured as total displacement across seven images acquired over 18 minutes total (yielding an inter-image interval of three minutes). This protocol preserves the typical throughput of our system while allowing multiple timepoint acquisitions per day of movement across several minutes.

When we simulated this protocol, it displayed marked improvement in measuring movement and age-related decline. Individual trajectories of decline were more qualitatively similar to those from minute-to-minute movement measurements compared to those estimated by the other protocols (cf. Fig 5H and Fig 5B). Additionally, movement measurements using this protocol were correlated with total movement measured from 1 fpm imaging over the total 3-hour intertimepoint interval (mean R^2^ = 0.44; Fig 5I-J). Most importantly, we found that measurement of movement decline using this scheme was able to identify decline in just over half of animals that had showed decline at a resolution of 1 fpm (14/26). These findings suggest that a simple modification to movement measurement that allows for frequent sampling of moderate-quality measurements may be sufficiently sensitive to capture early-life movement decline.

## Discussion

In this study, we questioned whether young adult individuals are subject to age-related declines in movement analogous to those seen in older individuals. Contrary to other work in this area, we found evidence that movement activity begins to decline as early as the first days of adulthood. Moreover, early-adult movement behavior is highly variable across individuals; in fact, the level of variation in total movement on this first day of adulthood is comparable to or higher than the variation typically seen in other life traits in wild-type *C. elegans*, including lifespan[11] and fecundity[38]. Taken together, these findings suggest that the vigorous movement that characterizes early adult life is not some generic state maintained consistently over time and across all individuals until some qualitative failure point. Rather, it is a relative and variable trait that declines smoothly over time, even in otherwise “healthy” animals.

The evidence that individuals show such variability in early-life movement contrasts with previous work from our group and others. In many studies, average movement has a nearly constant value representing “healthy” young-adult movement that only declines starting at days 6-8. We suspect that this high level of activity reflects a ceiling effect of the movement measurements used. That is, our results suggest that measurements based on brief intervals of observed activity may not have the sensitivity to detect inter- and intraindividual differences in movement during the first several days of adulthood. Indeed, a recent study of interindividual variability in movement during development using high-temporal resolution (3 fps) also suggests significant variation in movement activity at least on the first day of adulthood[39].

Motivated by these findings, we identified an intermediate measurement regime that provides a substantial improvement in sensitivity to early-adult movement. In particular, we find that the advantage of sampling movement more often over the day more than outweighs the disadvantages of conducting that sampling at a coarser time-scale (on the order of minutes). We hope that these results will inspire others to determine whether optimized movement measurement protocols in different culture systems can also be as or more accurate than current protocols while also requiring fewer images to be analyzed overall.

Several unique features of our culture system may limit the direct generalizability of these results to all settings. First, the “worm corral” system involves a relatively confined environment (a roughly circular bacterial food pad ≈1.5-1.6 mm, or ~1.5-1.75 typical body lengths, in diameter). This smaller area might influence behavior, as it is known animals on food tend to be more quiescent than those off food[37], potentially leading to underestimation of an animal’s overall capacity to move[14,37]. Moreover, the small food pad size places a ceiling over the maximum displacement measurable between timepoints, as total movement distance on the scale of minutes or more is often larger than the size of the pad. This is consistent with our observations that inter-imaging intervals greater than 5-10 minutes compromise movement measurements. In culture systems with larger movement arenas, longer inter-imaging intervals may be as or more useful than those employed here. Another feature of our culture system is the use of relatively high magnification compared to other long-term imaging apparatuses, which requires moving the stage to and focusing on individual animals one after another. In this context, our hybrid protocol provides the advantage that imaging bouts for one individual can be interleaved with imaging for others, increasing overall throughput. Optimal protocols may differ for other longitudinal observation systems in which many identifiable individuals can be simultaneously observed at once[12,40]. Last, we examined unstimulated movement only, which may be less sensitive for observing age-related changes across life[12,32].

Nevertheless, our finding of modest early-life declines in movement ability and/or propensity in most individuals provides a similar opportunity for researchers with different systems to calibrate and optimize their own motion-measurement protocols. Likewise, we suspect that the basic scheme of the optimized “hybrid” protocol we identified will be of use to others: the combination of moderately sensitive measurements repeated multiple times daily may be both more efficacious than infrequent, high-sensitivity imaging bouts and also allow greater throughput than continuous high-sensitivity imaging.

## Acknowledgements

The BA671[*spe-9(hc88)*] strain was provided by the CGC, which is funded by NIH Office of Research Infrastructure Programs (P40 OD010440). The authors appreciate critical feedback on early drafts of this manuscript from Drs. Marilyn Piccirillo and Arnav Moudgil.

---

**S1 Fig: Correlation between second-scale and minute-scale total movement over 15 minute intervals.** Comparison of total displacement at 1 fps over 15 minutes and the summed net displacement at 1 fpm over the same 15-minute interval (n = 48 animals) using the pilot 1 fps dataset. The overall correlation between total and net summed displacement is R^2^ = 0.81 (95% CI 0.67-0.90 generated by bootstrap resampling the measurements of random subsets of 43 animals/90% of animals).

**S2 Fig: Performance of Twice-Daily Video assessment of movement.** (A) The same individual in Fig 5A and 5E’s activity across all 6 days of life using a simulated twice-daily video measurement. (B) The correlation between the total displacement during the sampling period and the subsequent time (i.e. given by the intertimepoint interval). (C) Kernel density plot showing the distribution of correlations obtained for all individuals with the various intertimepoint interval

## Notes

### Competing Interest Statement

The authors have declared no competing interest.

